# Opposing roles of GSDMD and NINJ1 shape early IL-1β–dependent host defense during *Toxoplasma gondii infection*

**DOI:** 10.64898/2026.05.28.728484

**Authors:** Rafael Queiroz de Souza, Amanda Verdicchio Caniato, Luiza Zainotti Miguel Fahur Bottino, Mariana de Jesus Dias, Karina Ramalho Bortoluci

## Abstract

Inflammasome activation is a central component of host defense against *Toxoplasma gondii*, yet how downstream pyroptotic effectors shape infection outcomes remains incompletely understood. Here, we show that infection of primary macrophages with virulent type I *T. gondii* induces time-dependent NLRP3 inflammasome activation, caspase-1 cleavage, IL-1β secretion, and cell death. Unexpectedly, although caspase-1/11 was required for parasite control, the pore-forming effector gasdermin D (GSDMD) limited host resistance. GSDMD-deficient macrophages and mice exhibited enhanced control of parasite replication, despite reduced cell death. Mechanistically, GSDMD deficiency uncoupled inflammasome activation from pyroptotic lysis, leading to accelerated and increased early IL-1β release. Exogenous IL-1β restored parasite control in wild-type macrophages, indicating that early cytokine availability is a key determinant of resistance. Enhanced control in GSDMD-deficient cells was associated with increased NLRP3 inflammasome assembly and was partially reversed by pharmacological inhibition of NLRP3, supporting a role for early inflammasome signaling in parasite restriction. In contrast, the membrane rupture mediator NINJ1 promoted host protection. NINJ1-deficient macrophages displayed impaired IL-1β release and increased parasite replication, indicating that terminal cell lysis and cytokine release can be functionally dissociated. Pharmacological inhibition of GSDMD using disulfiram recapitulated the protective phenotype in both murine and human macrophages and reduced parasite burden *in vivo*, without directly affecting parasite viability. Together, these findings reveal that inflammasome-dependent cytokine production and pyroptotic cell death exert distinct and, in some cases, opposing roles during *T. gondii* infection. While NINJ1 supports protective responses, GSDMD-driven pyroptosis constrains early IL-1β–mediated immunity. Targeting GSDMD may therefore represent a therapeutic strategy to enhance host control of intracellular pathogens.

## Introduction

*Toxoplasma gondii* is an obligatory intracellular protozoan that establishes lifelong infection in a broad range of warm-blooded hosts. Approximately one-third of the global population is chronically infected, reflecting the parasite’s remarkable ability to evade and modulate host immunity while preventing sterilizing clearance (1,2).

Host recognition of *T. gondii* relies on coordinated innate immune pathways, including Toll-like receptors, interferon-inducible GTPases, and inflammasomes (3,4). Among these, inflammasome signaling is central to host resistance. Infection activates multiple inflammasome sensors in both human and murine cells, culminating in caspase-1 activation and IL-1β release (5–10). Consistently, genetic ablation of key inflammasome components, such as NLRP3, NLRP1, ASC, caspase-1/11, IL-1R, and IL-18, results in increased parasite burden and reduced survival *in vivo* (11–15), underscoring the importance of this pathway in anti-parasitic defense.

While upstream inflammasome sensors have been extensively characterized during *T. gondii* infection, considerably less is known about the downstream pore-forming effector Gasdermin D (GSDMD). In most infectious settings, GSDMD is considered as protective, promoting pathogen clearance through pyroptosis and cytokine release (16–20). However, emerging evidence indicates the role of GSDMD may be context-dependent and vary according to pathogen and cellular compartment (21). In the setting of *T. gondii* infection, GSDMD has been implicated in neutrophil extracellular trap formation (22) and microglial IL-1α release (23), yet its role in macrophages, the primary niche for parasite replication, remains undefined.

Here, we demonstrate that GSDMD exerts an unexpected regulatory function during acute infection with the RH strain of *T. gondii*. In contrast to its protective roles described in other infectious contexts, GSDMD deficiency reduced parasite burden both *in vitro* and *in vivo*. Mechanistically, absence of GSDMD enhanced early NLRP3 inflammasome activation and IL-1β production, resulting in improved parasite control in a NLRP3-dependent manner. Collectively, these findings challenge the current view of GSDMD as uniformly protective and identify a previously unrecognized role in macrophage immunity against *T. gondii*.

## Materials and Methods

### Animals

This study was reviewed and approved by the Ethics Committee on the Use of Animals of the Federal University of São Paulo (UNIFESP) and is registered under protocol number 9154060923. C57BL/6, Caspase-1/11^⁻/⁻^, GSDMD^⁻/⁻^ mice (kindly provided by Dr. Petr Broz, University of Lausanne, and supplied by Prof. Sergio Costa Oliveira, University of São Paulo) and NINJ1^⁻/⁻^ mice (originally obtained from Genentech and kindly provided by Prof. Dario Simões Zamboni, Faculty of Medicine University of São Paulo, Ribeirão Preto Campus) aged 6–10 weeks, were housed at the Center for the Development of Experimental Models for Medicine and Biology (CEDEME, UNIFESP) under specific pathogen-free (SPF) conditions, in microisolator cages with free access to food and water.

### Parasites

For the experiments, the Type I RH strain of *Toxoplasma gondii* genetically modified to express yellow fluorescent protein (*T. gondii* RH YFP) was used. Parasites were stored in liquid nitrogen and, upon thawing, were maintained in cultures of immortalized human foreskin fibroblasts (Hs27), grown in DMEM supplemented with 5% fetal bovine serum (FBS) at 37 °C and 5% CO₂. After host cell lysis resulting from parasite replication, the supernatant containing *T. gondii* tachyzoites was collected and centrifuged (900 × g, 4 °C, 10 min). The resulting pellet was resuspended in RPMI supplemented with 3% FBS for immunofluorescence assays or in Opti-MEM for Western blot analyses. Parasite viability was assessed using Trypan blue exclusion in a Neubauer hemocytometer, and parasites were adjusted to a final concentration corresponding to a multiplicity of infection (MOI) of 2 parasites per cell. The parasite strain was kindly provided by Dr. Ricardo Tostes Gazzinelli (UFMG/Fiocruz – René Rachou/UMass, USA).

### Bone marrow–derived macrophage differentiation

Bone marrow cells were obtained from femur and tibia flushes using sterile cold PBS. Cells were centrifuged (500 × g, 4 °C, 10 min), treated with hemolytic buffer for 2 min, washed with sterile PBS, centrifuged again (500 × g, 4 °C, 10 min), and counted using a Neubauer hemocytometer. After counting, 4.5 × 10⁶ cells were seeded in non-adherent 75 cm² flasks containing R10% medium [RPMI 1640 (Gibco) supplemented with 10% FBS (Gibco), 1% penicillin/streptomycin (Gibco), 1% GlutaMAX (Gibco), 1% HEPES, 1% sodium pyruvate (Gibco), 1% non-essential amino acids (Gibco), 1% vitamins, 20% L929 cell-conditioned supernatant, and 55 µM β-mercaptoethanol (Sigma-Aldrich)], and incubated at 37 °C and 5% CO₂. Culture medium was completely replaced on days 3 and 5. On day 7, the cell supernatant was discarded, sterile PBS containing 2% FBS (Gibco) and 2 mM EDTA (Merck) was added, and cells were maintained on ice for 15 min. Bone marrow–derived macrophages (BMDMs) were then plated and incubated for 18 h at 37 °C and 5% CO₂ prior to infection or stimulation on the following day.

### BMDM infection and stimulation

BMDMs, treated or not with LPS (200 ng/mL) for 3 h at 37 °C and 5% CO₂, were infected with the *T. gondii* RH YFP strain at a ratio of 2:1 (parasite:cell). Parasites were maintained at 37 °C and 5% CO₂ throughout the infection period. When required, cells were treated with the NLRP3 inflammasome inhibitor MCC950 (10 µM) or the GSDMD inhibitor disulfiram (10 µM) for 1 h at 37 °C and 5% CO₂ prior to infection, and the inhibitors were maintained throughout the experiment. When indicated, 4 × 10⁵ *T. gondii* tachyzoites were exposed to UV light for 15 min or incubated at 100 °C for 5 min in a thermoblock prior to infection. As a positive control for inflammasome activation, cells were primed with LPS (200 ng/mL) (InvivoGen, San Diego, CA, USA) for 3 h and subsequently treated with the agonist nigericin (10 µM) (InvivoGen) for 60 min. All infections and stimuli were performed in R3% medium, except for Western blot assays, which were conducted using Opti-MEM (Gibco) supplemented with 0.029 mM sodium bicarbonate (Sigma-Aldrich), 0.23 mM streptomycin (Sigma-Aldrich), and 0.25 mM penicillin (Sigma-Aldrich), pH 7.1.

### *In vivo* infection and parasite load quantification

Mice were infected with the *T. gondii* RH YFP strain by intraperitoneal injection of 1 × 10⁴ tachyzoites diluted in 200 µL of sterile PBS. Spleens were aseptically collected and immediately transferred to RPMI-1640 supplemented with 3% fetal bovine serum (FBS). Organs were mechanically dissociated by passage through a 100 µm cell strainer using a sterile syringe plunger. The resulting cell suspension was carefully homogenized, and extracellular parasites present in the suspension were quantified using a Neubauer hemocytometer. For *in vivo* experiments using the GSDMD inhibitor, disulfiram was diluted in a solution containing 10% DMSO, 2% Tween-20, and 88% corn oil to achieve a final concentration of 50 mg/kg. Animals were treated daily, receiving an injection of 100 µL of the prepared solution. Alternatively, peritoneal cells from infected mice were plated in R3% medium [RPMI 1640 (Gibco) supplemented with 3% fetal bovine serum (FBS) and after incubation (*ex vivo*), cells were fixed for subsequent immunofluorescence analysis.

### *In vitro* infection quantification

All immunofluorescence experiments were performed in 96-well plates, with BMDMs seeded at a density of 2 × 10⁵ cells per well. To assess the infection index, the IN Cell Analyzer 2200 system (GE Healthcare), which enables automated image acquisition, was used. Briefly, after the infection period, the supernatant was collected and reserved. Plates were fixed at room temperature for at least 15 min using 3% paraformaldehyde (Sigma-Aldrich) diluted in PBS. Wells were then washed with warm PBS, incubated with DAPI at 5 µg/mL (Sigma-Aldrich) diluted in PBS for 2 min, and subsequently replaced with PBS. A total of 9–12 images per well were acquired using the IN Cell Analyzer 2200 and analyzed using ImageJ software. Infection was quantified using two complementary approaches: (i) the percentage of infected cells, calculated as the number of YFP-positive cells relative to the total number of DAPI-positive cells, and (ii) the parasite burden per macrophage, determined by measuring the total YFP-positive area (µm²) per field and normalizing it to the number of macrophages, allowing discrimination between differences in infection prevalence and intracellular parasite load.

### ELISA-detected inflammasome activation in BMDMs

BMDMs were seeded in 96-well plates at a density of 2 × 10⁵ cells per well and infected at a multiplicity of infection (MOI) of 2:1. IL-1β production in culture supernatants was quantified by ELISA using a commercial kit from Thermo Fisher Scientific (Invitrogen; Cat# 88-7013-88). The levels of cleaved inflammasome effectors in the supernatant were determined using ELISA kits from AdipoGen. Cleaved gasdermin D (GSDMD) was quantified using a kit specific for the C-terminal fragment of GSDMD (AdipoGen, Cat# AG-45B-0011-KI01), while active caspase-1 was measured using a kit that detects the enzymatically active p20 subunit of caspase-1 (AdipoGen, Cat# AG-46B-0003-KI01). All assays were performed according to the manufacturers instructions. The absorbance of the plate was read at 450 nm on a 165 SpectraMax instrument.

### Cell viability

Cell viability was assessed by measuring the release of Lactate Dehydrogenase (LDH) using a commercial LDH Cytotoxicity Assay Kit (Invitrogen, Cat# C20301), according to the manufacturer’s instructions. LDH activity was quantified in the culture supernatants collected after the indicated stimulation or infection times.

### Immunofluorescence staining

Immunofluorescence assays for ASC and NLRP3 were performed in black, clear-bottom 96-well plates (Greiner) suitable for high-content imaging. Cells were plated under standard conditions and incubated for 18 h at 37 °C and 5% CO₂. The following day, cells were infected with *Toxoplasma gondii* at a multiplicity of infection (MOI) of 2:1 (parasite:cell) for 2, 4, or 18 h, or treated with the positive control nigericin for 1 h, at 37 °C and 5% CO₂. After stimulation or infection, culture supernatants were removed and cells were fixed for at least 15 min with 3% paraformaldehyde (Sigma-Aldrich) diluted in PBS at room temperature. Cells were then washed with PBS and incubated for 30 min at room temperature with a blocking/permeabilization solution containing 10% bovine serum albumin (BSA; Sigma-Aldrich), 1% fetal bovine serum (FBS; LGC), and 0.5% Triton X-100 (Sigma-Aldrich) diluted in PBS. Following one gentle wash with warm PBS, cells were incubated overnight (18 h) at 4 °C with either anti-ASC antibody (1:1000; Santa Cruz Biotechnology, sc-514414) or anti-NLRP3 antibody (1:500; AdipoGen, AG-20B-0014-C100). Cells were subsequently washed twice with PBS and incubated with Alexa Fluor 647–conjugated secondary antibody (1:1000; Invitrogen, A21235) for 1 h at room temperature, protected from light. After additional washes, nuclei were stained with DAPI (5 µg/mL; Sigma-Aldrich) for 2 min, followed by replacement with PBS. Images were acquired using an IN Cell Analyzer 2200 (GE Healthcare), and image analysis was performed using ImageJ software.

### Western blotting

For western blotting experiments, cells were plated in technical duplicates for each condition at a density of 1 × 10⁶ cells per well in 24-well plates, using R10% medium, and incubated for 18 h at 37 °C and 5% CO₂. After incubation, the culture medium was replaced with Opti-MEM (Gibco). For infection assays, macrophages were primed or not with LPS (200 ng/mL) for 3 h at 37 °C and 5% CO₂ and subsequently infected with *Toxoplasma gondii* RH YFP strain at a multiplicity of infection (MOI) of 2:1 (parasite:cell) for 18 h. Following infection, culture supernatants were collected and proteins were precipitated using a methanol/chloroform protocol. Adherent cells were lysed using RIPA buffer (Sigma-Aldrich) supplemented with protease inhibitor cocktail (1:1000). Technical duplicates were pooled, resulting in one combined protein sample per condition. Prior to electrophoresis, samples were mixed with sample buffer (Thermo Fisher Scientific) containing 710 mM β-mercaptoethanol (Sigma-Aldrich) and denatured. Proteins were separated by SDS-PAGE using 13.5% polyacrylamide gels for 90 min and subsequently transferred to PVDF membranes (Merck) using a wet transfer system at 100 V for 1 h. Membranes were blocked for 1 h at room temperature with 5% bovine serum albumin (BSA; Sigma-Aldrich) diluted in TBS-T (0.05% Tween-20), washed three times with TBS-T, and incubated overnight at 4 °C under agitation with the following primary antibodies: anti–caspase-1 (1:500; AdipoGen), anti-GSDMD (1:1000; Abcam), anti–IL-1β (1:500; Sigma-Aldrich), anti-NLRP3 (1:500; AdipoGen), and anti–β-actin (1:3000; Sigma-Aldrich). After incubation, membranes were washed three times with TBS-T and incubated with appropriate HRP-conjugated secondary antibodies for 1 h at room temperature. Membranes were then washed again and protein detection was performed by chemiluminescence using ECL substrate (Santa Cruz Biotechnology). Signals were acquired using the Alliance 4.7 software on the Uvitec Cambridge imaging system. When necessary, membranes were stripped using ReBlot commercial stripping solution (Merck) for 30 min at room temperature under agitation and reprobed.

### THP-1 cell culture and macrophage-like differentiation

THP-1 monocytic cells were cultured in RPMI 1640 medium supplemented with 10% heat-inactivated fetal bovine serum (FBS), 1% L-glutamine, and 1% penicillin–streptomycin (Invitrogen) and maintained at 37 °C in a humidified atmosphere containing 5% CO₂. For differentiation into macrophage-like cells, THP-1 cells were treated with phorbol-12-myristate-13-acetate (PMA) at a final concentration of 10 nM for 24 h. After differentiation, cells were allowed to rest for an additional 24 h in RPMI 1640 medium without PMA prior to experimental procedures.

### Data analysis

All statistical analyses were performed using GraphPad Prism software version 9.3.0 (GraphPad Software Inc.), applying the most appropriate statistical test for each dataset.

## Results

### *T. gondii* triggers inflammasome activation in primary macrophages in a time-dependent manner

Macrophages are key sentinels of *Toxoplasma gondii* infection and play a central role in initiating innate immune responses required for parasite control (24,25). Previous studies have shown that *T. gondii* proteins can modulate inflammasome signaling in host cells, acting either as activators or suppressors of inflammasome pathways (26–31). However, whether infection with highly virulent *T. gondii* strains directly triggers inflammasome activation in primary macrophages remains incompletely defined. To address this, we infected bone marrow-derived macrophages (BMDMs) with YFP-expressing tachyzoites from the highly virulent type I RH strain at a multiplicity of infection (MOI) of 2.

First, we observed that *T. gondii* infection induced the expression of NLRP3 (Figure 1A) and pro-IL-1β (Figure 1B) in BMDMs in a time-dependent manner. Quantification of intracellular parasite burden revealed a progressive increase in replication over time, indicating active intracellular proliferation (Figure 1C). Notably, *T. gondii* infection promoted time-dependent ASC speck formation (Figure 1D), cleaved caspase-1 (Figure 1E), mature IL-1β (Figure 1F), and GSDMD C-terminal (Figure 1G), together with increased lactate dehydrogenase (LDH) release in culture supernatants (Figure 1H), indicating the induction of inflammasome activation.

**Figure 1.**
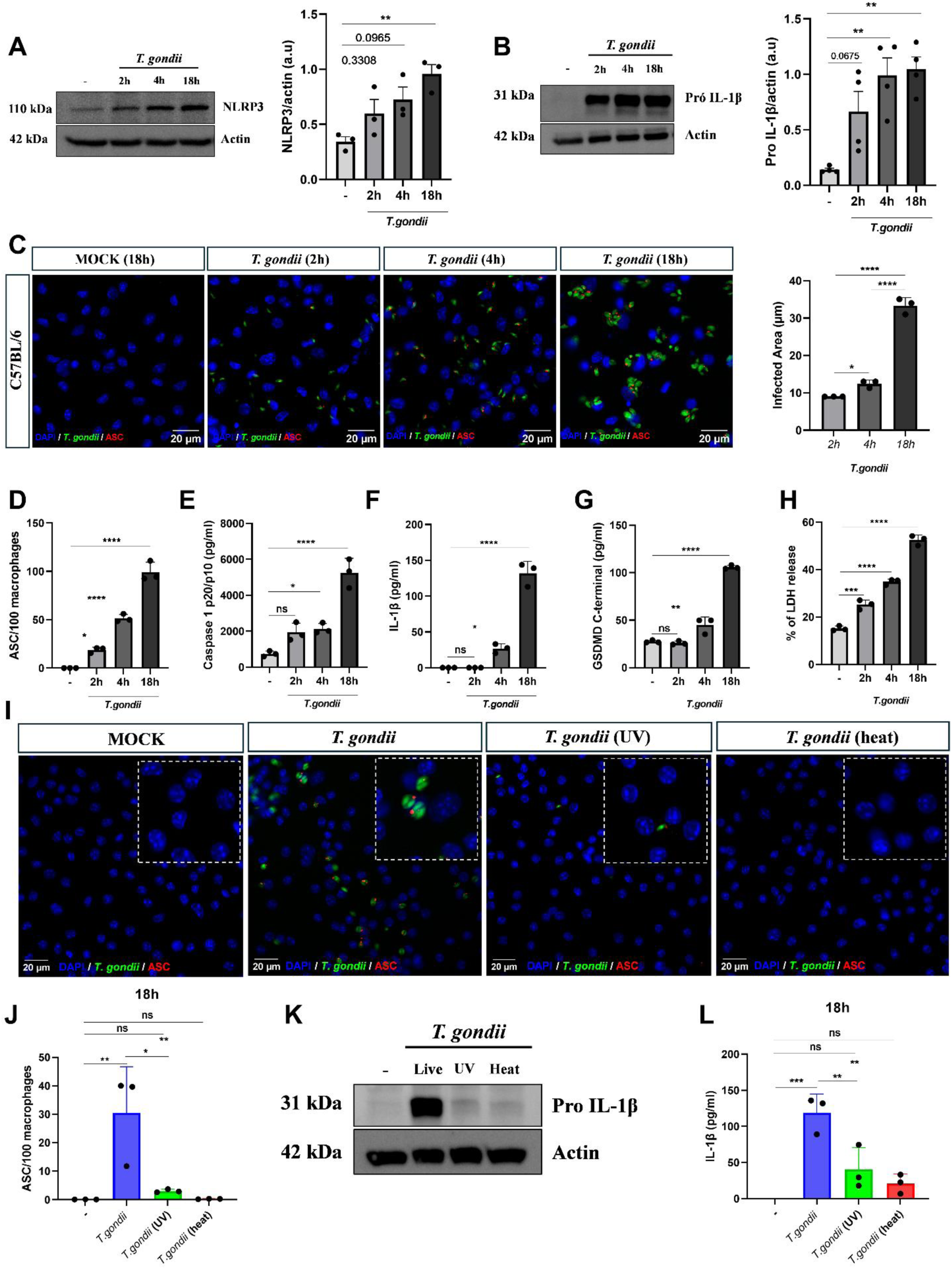
*T. gondii* triggers inflammasome activation in primary macrophages in a time-dependent manner. Bone marrow–derived macrophages (BMDMs) from C57BL/6 mice were infected with *T. gondii* RH-YFP at a multiplicity of infection (MOI) of 2:1 for the indicated times. (A–B) BMDMs (1 × 10⁶ cells) were infected for 2, 4, or 18 h and cell lysates were analyzed by Western blot to assess inflammasome-related proteins. β-actin was used as a loading control. Representative blots and quantification from independent experiments are shown. (C–D) BMDMs (2 × 10⁵ cells/well) were infected and subsequently fixed and stained with DAPI. Parasite burden (C) and ASC speck formation (D) were quantified by immunofluorescence microscopy. (E) Caspase-1 (F) IL-1β secretion and (G) GSDMD cleavage were quantified in culture supernatants by ELISA. (H) Cell death was evaluated by LDH release in culture supernatants. All graphs present data from one representative experiment of at least three independent experiments performed in technical triplicates. (I–J) BMDMs were infected with viable or non-viable *T. gondii* RH-YFP (MOI 2:1) for 18 h. Representative immunofluorescence images (I) and quantification of ASC puncta (J) are shown. Data represents the mean of three independent experiments performed in technical triplicates. (K) BMDMs were infected for 18h and analyzed by Western blot following cell lysis in RIPA buffer. β-actin was used as a loading control. Representative blot of two independent experiments. (L) IL-1β release in culture supernatants was measured by ELISA. Images were acquired using an IN-Cell Analyzer 2200 microscope and quantified using ImageJ. Data represents the mean of three independent experiments performed in technical triplicates. Statistical analysis was performed using one-way ANOVA or Student’s t-test. *p < 0.05, **p < 0.01, ***p < 0.001, ****p < 0.0001; ns, not significant.

To determine whether inflammasome activation depends on parasite viability and replication, RH tachyzoites were inactivated by ultraviolet (UV) irradiation or heat killing prior to infection. BMDMs infected with viable parasites exhibited robust ASC speck formation, whereas UV-irradiated or heat-killed tachyzoites failed to induce inflammasome assembly (Figure 1I-J). Consistently, only viable parasites promote pro-IL-1β expression (Figure 1K) and IL-1β release (Figure 1L). Together, these findings demonstrate that *T. gondii* replication promotes the induction of inflammasome components, inflammasome assembly and canonical activation, culminating in pyroptotic cell death.

### GSDMD limits macrophage-mediated control of *Toxoplasma gondii*

The protective role of the inflammasome and caspase-1/11 during *Toxoplasma gondii* infection has been well established, particularly in models using type II parasite strains (12). However, how these pathways operate during infection with highly virulent type I strains remains less clear, and the specific contribution of the downstream effector GSDMD has not been defined. As expected, Casp1/11^⁻/⁻^ macrophages were more susceptible to infection than wild-type (WT) cells (Supplemental Figure 1A), exhibiting a higher percentage of infected cells (Supplemental Figure 1B) and a larger infected area (Supplemental Figure 1C). Consistently, Casp1/11^⁻/⁻^ mice displayed increased parasite burden in the spleen compared with C57BL/6 controls (Supplemental Figure 1D), confirming a critical role for caspase-1/11 in restricting *T. gondii* replication.

Unexpectedly, GSDMD^−/-^ BMDMs showed significantly improved control of parasite replication at 18 h post-infection (Figure 2A). This phenotype was associated with a decreased YFP⁺/DAPI⁺ area ratio compared with WT macrophages (Figure 2B) and a reduced percentage of infected cells (Figure 2C). As expected, GSDMD-deficient macrophages were less susceptible to parasite-induced cell death, as indicated by reduced LDH release relative to WT cells (Figure 2D). Notably, no differences parasite burden between WT and GSDMD^−/-^ BMDMs were observed at early time points (2 or 4 h post-infection; Figure 2A–B), suggesting that GSDMD deficiency does not affect parasite invasion but instead influences host–parasite interactions at later stages of infection.

**Figure 2.**
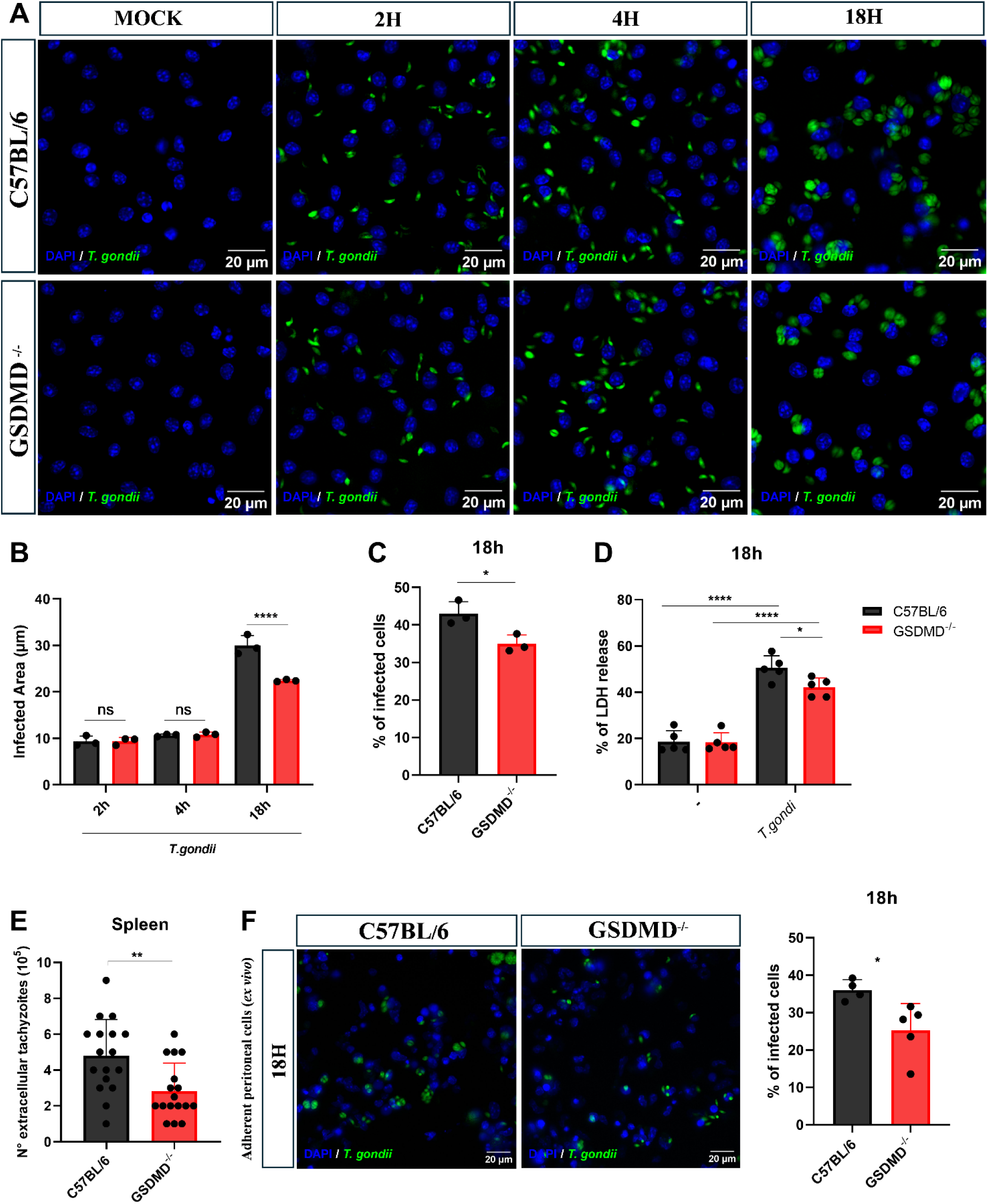
GSDMD limits macrophage-mediated control of *T. gondii*. Bone marrow–derived macrophages (BMDMs; 2 × 10⁵ cells/well) from C57BL/6 (WT) and GSDMD^⁻/⁻^ mice were infected with *T. gondii* RH-YFP tachyzoites at a multiplicity of infection (MOI) of 2 for the indicated times (2, 4, or 18 h). After infection, cell culture supernatants were collected and cells were fixed with PFA and stained with DAPI. (A) Representative immunofluorescence images of infected macrophages. (B) Parasite burden was quantified as the ratio between the YFP⁺ parasite area and the number of macrophages. (C) The percentage of infected cells was determined from immunofluorescence images and used to calculate the infection index. (D) Cell death was evaluated by measuring LDH release in culture supernatants. Alternatively, C57BL/6 and GSDMD^⁻/⁻^ mice (6–10 weeks old) were infected intraperitoneally with 1 × 10⁴ *T. gondii* RH tachyzoites. (E) Four days post-infection, animals were euthanized and extracellular tachyzoites in the spleen were quantified. (F) Peritoneal cells were recovered by lavage four days post-infection, plated (5 × 10⁵ cells/well), and incubated for 18 h. Non-adherent cells were removed and adherent cells were fixed with PFA and stained with DAPI. Representative images are shown and infection was quantified in adherent cells. Data in panel B presents data from one representative experiment of at least three independent experiments performed in technical triplicates. Panel C represents the mean of three independent experiments performed in technical triplicates. Panel D represents five independent experiments. Panel E shows pooled data from four independent experiments (WT n = 17; GSDMD^⁻/⁻^ n = 17), with each dot representing one animal. In panel F, each data point represents one individual animal (WT n = 4; GSDMD^⁻/⁻^ n = 5), calculated as the mean of five technical replicates from the same animal. Statistical analysis was performed using Student’s *t*-test or two-way ANOVA. *p* < 0.05, p < 0.01, *p* < 0.001, p < 0.0001; ns, not significant.

Consistent with these *in vitro* findings, GSDMD^⁻/⁻^ mice displayed significantly reduced parasite burden in the spleen compared with C57BL/6 controls (Fig. 2E), indicating enhanced resistance to *T. gondii* replication in the absence of GSDMD. Similarly, adherent peritoneal cells isolated from infected GSDMD^⁻/⁻^ mice showed a lower percentage of infected cells relative to cells obtained from WT animals (Fig. 2F), further supporting a resistant phenotype.

Together, these findings reveal an unexpected detrimental role for GSDMD during *T. gondii* infection, indicating that GSDMD activation limits effective macrophage-mediated control of parasite replication.

### NINJ1 promotes macrophage-mediated control of *T. gondii*

Although GSDMD activation is indispensable for the execution of pyroptosis, it is not solely responsible for the terminal rupture of the plasma membrane during this inflammatory form of cell death. Recent studies have demonstrated that the final lytic event constitutes a distinct process regulated by the membrane protein NINJ1 (32). Given the complementary roles of GSDMD and NINJ1 during pyroptosis, we sought to investigate the role of NINJ1 in this infectious context to determine whether these pyroptosis-associated molecules exert redundant or distinct functions in response to *T. gondii* infection.

As expected, LDH release in response to nigericin treatment was completely dependent on NINJ1 expression (Figure 3A). In contrast, IL-1β release was not impaired in the absence of this molecule (Figure 3B), indicating that although NINJ1 is essential for plasma membrane rupture and terminal cell lysis, it is dispensable for canonical inflammasome activation and cytokine processing.

**Figure 3.**
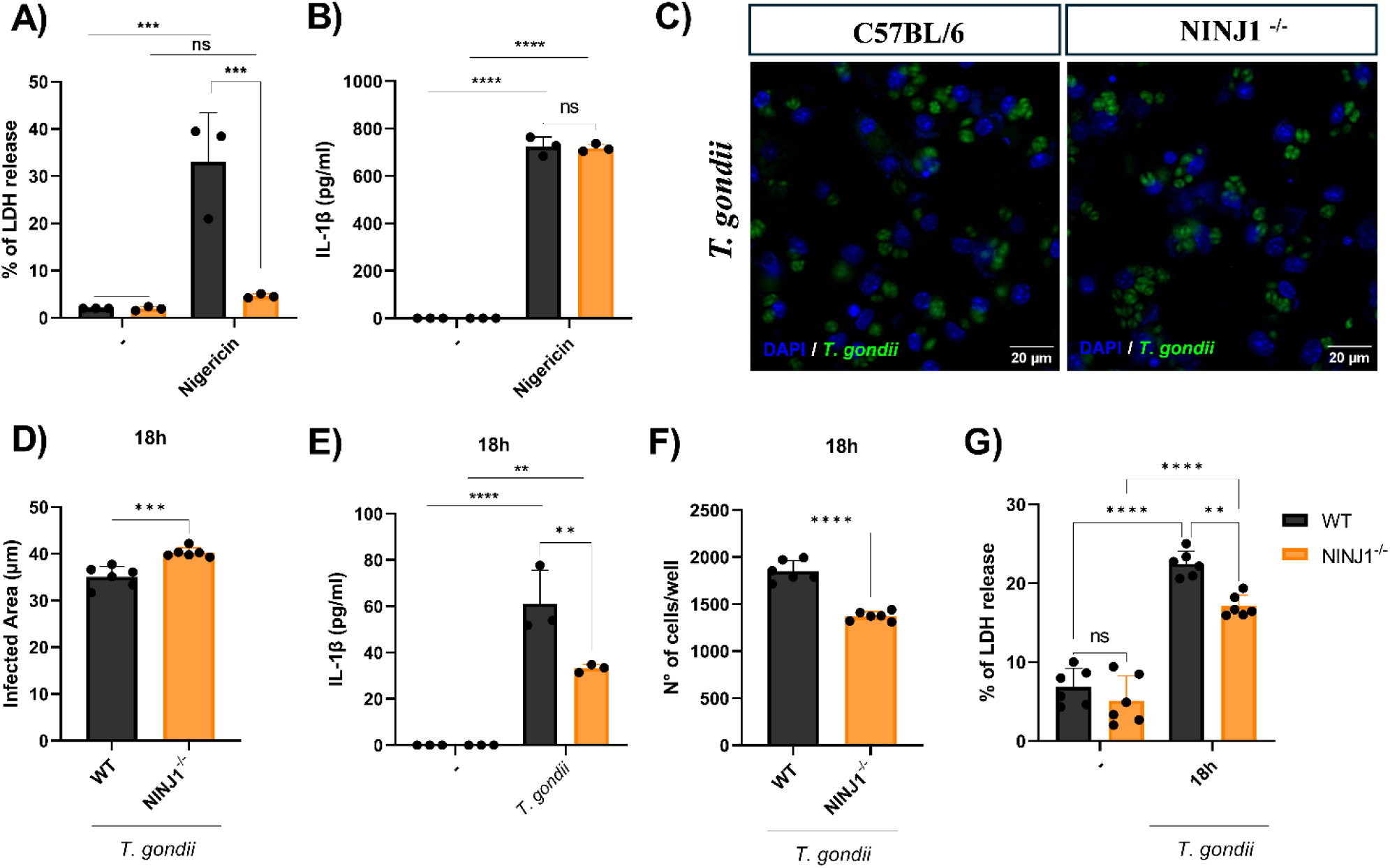
NINJ1 promotes macrophage-mediated control of *T. gondii*. (A and B) Bone marrow–derived macrophages (BMDMs; 2 × 10⁵ cells/well) from C57BL/6 (WT) and NINJ1^⁻/⁻^ were primed with LPS (200 ng/mL) for 3 h and subsequently stimulated with nigericin (10 µM) for 1 h as a positive control for inflammasome activation. (C-G) Alternatively, Bone marrow–derived macrophages (BMDMs; 2 × 10⁵ cells/well) from C57BL/6 (WT) and NINJ1^⁻/⁻^ mice were infected with *T. gondii* RH-YFP tachyzoites at a multiplicity of infection (MOI) of 2 for 18 h. After infection, cell culture supernatants were collected and cells were fixed with PFA and stained with DAPI. (A) Cell death was evaluated by measuring LDH release in culture supernatants. (B) IL-1β secretion was measured in culture supernatants by ELISA. (C) Representative immunofluorescence images of infected macrophages. (D) Parasite burden was quantified as the ratio between the YFP⁺ parasite area and the number of macrophages. (E) IL-1β secretion was measured in culture supernatants by ELISA. (F) The total number of cells remaining adherent in each well was quantified from the immunofluorescence images using ImageJ. (G) Cell death was evaluated by measuring LDH release in culture supernatants. The data shown are representative of three independent experiments performed with at least technical triplicates, except for panels A, F, and G, which are representative of two independent experiments. Statistical analysis was performed using Student’s *t*-test or two-way ANOVA. *p* < 0.05, p < 0.01, *p* < 0.001, p < 0.0001; ns, not significant.

Interestingly, whereas GSDMD deficiency was associated with enhanced parasite control, NINJ1^⁻/⁻^ macrophages exhibited higher levels of *T. gondii* replication relative to WT BMDMs (Figure 3C-D). The increased susceptibility to *T. gondii* infection observed in NINJ1-deficient macrophages was associated with reduced IL-1β release (Fig. 3E), a lower number of viable cells remaining in the culture after 18 h of infection (Fig. 3F) and decreased LDH secretion (Fig. 3G) compared to WT BMDMs. Overall, these results support a protective role for NINJ1 in *T. gondii* infection and highlight that NINJ1 and GSDMD mediate distinct functions in this context.

### GSDMD deficiency leads to an accelerated IL-1β response during *T. gondii* infection

To understand why GSDMD deficiency enhances resistance to *T. gondii*, we next examined inflammasome activation in GSDMD^−/-^ macrophages. First, as expected, a time-dependent GSDMD cleavage in response to *T. gondii* was detected in WT but not in GSDMD^⁻/⁻^ macrophages (Figure 4A).

**Figure 4.**
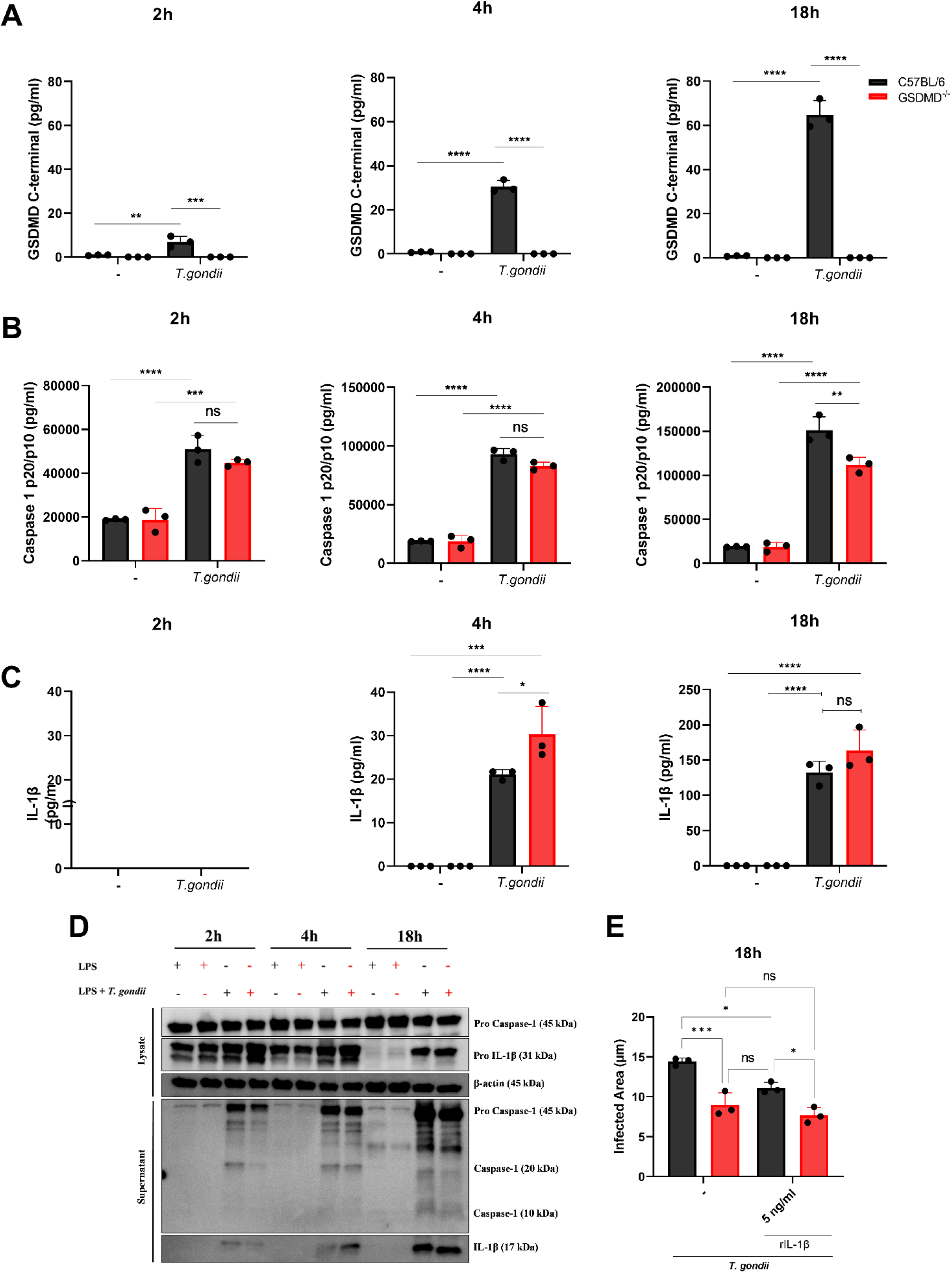
GSDMD deficiency leads to an accelerated IL-1β response during *T. gondii* infection. (A–C and E) Bone marrow–derived macrophages (BMDMs; 2 × 10⁵ cells/well) from C57BL/6 (WT) and GSDMD⁻/⁻ mice were infected with *T. gondii* RH-YFP tachyzoites at a multiplicity of infection (MOI) of 2 for 2, 4, or 18 h. After infection, cell culture supernatants were collected. (A) GSDMD cleavage in culture supernatants was quantified by ELISA. (B) Caspase-1 cleavage in culture supernatants was quantified by ELISA. (C) IL-1β secretion was measured in culture supernatants by ELISA. Graphs show data from one representative experiment of at least three independent experiments performed in technical triplicates. (D) Alternatively, BMDMs (1 × 10⁶ cells/well) from C57BL/6 and GSDMD⁻/⁻ mice were primed with LPS (200 ng/mL) for 3 h and infected with *T. gondii* RH-YFP tachyzoites (MOI 2) for 18 h. Supernatants were collected and precipitated, and cell lysates were prepared using RIPA buffer for protein detection in both fractions. β-actin was used as a loading control for normalization of protein levels in cell lysates. Data shown are representative of two independent experiments. (E) Parasite burden was quantified as the ratio between the YFP⁺ parasite area and the number of macrophages. Data shown are representative of one experiment. Statistical analysis was performed using two-way ANOVA. *p* < 0.05, p < 0.01, *p* < 0.001, p < 0.0001; ns, not significant.

In contrast to the results observed with nigericin (a classical NLRP3 agonist) (Supplemental Figure 2), cleaved caspase-1 was readily detectable in both *T. gondii*-infected WT and GSDMD^−/-^ macrophages at 2 and 4 hours post-infection, indicating that early caspase-1 activation occurs independently of GSDMD (Figures 4B and 4D). At 18 hours post-infection, cleaved caspase-1 levels increased in both groups, with higher levels observed in WT cells (Figures 4B and 4D).

Despite its dependence on caspase-1/11 (Supplemental Figure 1) and partial dependence on NINJ1 (Figure 3E), IL-1β release during *T. gondii* infection was preserved in GSDMD-deficient cells. In fact, GSDMD⁻/⁻ macrophages produced significantly higher levels of IL-1β at 4 hours post-infection compared to WT cells. By 18 hours, IL-1β levels were comparable between groups, although a trend toward increased secretion persisted in GSDMD-deficient macrophages (Figures 4C and 4D).

Given the link between IL-1β production and infection control in our model, we next asked whether exogenous IL-1β could modulate parasite replication. Recombinant IL-1β was added to macrophage cultures 4 hours post-infection, resulting in a reduction in parasite burden in WT cells to levels comparable to those observed in GSDMD⁻/⁻ macrophages (Figure 4E).

Together, these findings uncouple inflammasome-dependent cytokine release from GSDMD-mediated pyroptosis and reveal that GSDMD activation can restrain early IL-1β–driven host defense during *T. gondii* infection.

### Early NLRP3 inflammasome activation mediates enhanced resistance of GSDMD^−/-^ macrophages to *T. gondii* infection

Given the increased IL-1β release observed in GSDMD^⁻/⁻^ macrophages and previous evidence that NLRP3 inflammasome activation is required for effective control of *T. gondii* replication, we next examined NLRP3 inflammasome assembly during infection.

Immunofluorescence analysis revealed increased NLRP3 puncta formation in GSDMD^⁻/⁻^ BMDMs compared with WT cells at all time points analyzed (Figure 5A), consistent with the elevated IL-1β levels detected at early stages of infection (Figures 4C–D). Notably, the increased number of NLRP3 puncta was accompanied by a higher proportion of GSDMD^⁻/⁻^ macrophages remaining adherent during infection, whereas WT cells displayed progressive detachment (Figure 5B). These observations suggest that enhanced NLRP3 activation occurs in the context of improved cell viability.

**Figure 5.**
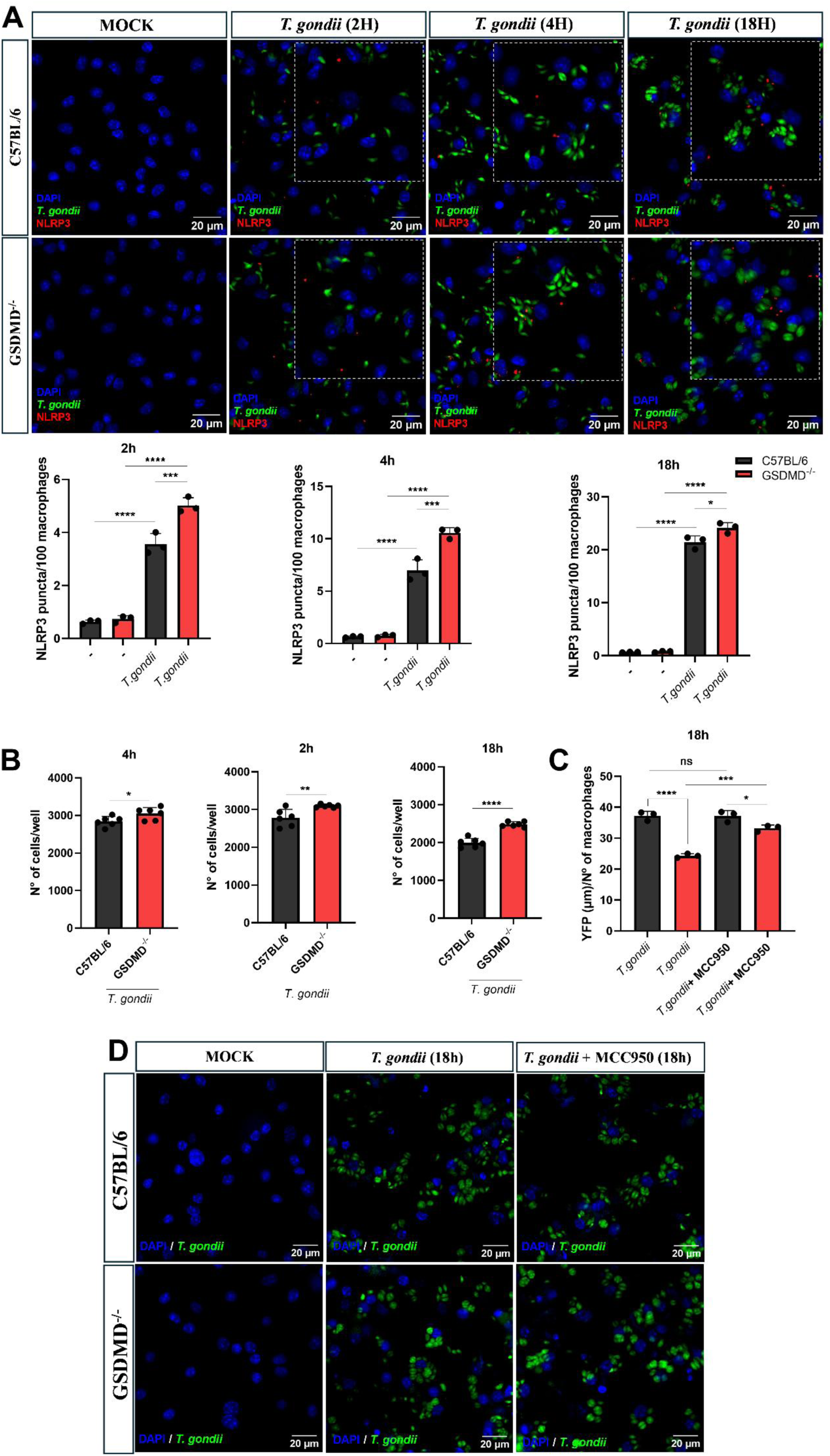
Early NLRP3 inflammasome activation mediates enhanced resistance of GSDMD⁻/⁻ macrophages to *T. gondii* infection. Bone marrow–derived macrophages (BMDMs; 2 × 10⁵ cells/well) from C57BL/6 (WT) and GSDMD⁻/⁻ mice were infected with *T. gondii* RH-YFP tachyzoites at a multiplicity of infection (MOI) of 2 for 2, 4, or 18 h. After infection, cell culture supernatants were collected and cells were fixed with PFA and stained with DAPI. (A) NLRP3 puncta formation was evaluated by immunofluorescence. Images were automatically acquired using an IN Cell Analyzer 2200 microscope and quantified using ImageJ. The graph shows data from one representative experiment of two independent experiments performed in technical triplicates. (B) The total number of cells remaining adherent in each well was quantified from the immunofluorescence images using ImageJ. Data are representative of three independent experiments. (C–D) Alternatively, BMDMs (2 × 10⁵ cells/well) from C57BL/6 and GSDMD^⁻/⁻^ mice were pretreated or not with the NLRP3 inhibitor MCC950 (10 µM) for 1 h prior to infection with *T. gondii* RH-YFP tachyzoites (MOI 2) for 18 h. Infection rates were determined by immunofluorescence microscopy using the IN Cell Analyzer 2200 system and quantified with ImageJ. Graphs show data from one representative experiment of at least four independent experiments.

Based on these findings, we hypothesized that the enhanced resistance to *T. gondii* replication observed in GSDMD^⁻/⁻^ macrophages results from increased NLRP3 inflammasome activation. To test this, macrophages were pretreated with MCC950, a selective NLRP3 inhibitor, prior to infection. While MCC950 had no significant effect on parasite replication in WT BMDMs, it partially reversed the resistant phenotype observed in GSDMD^⁻/⁻^ macrophages (Figure 5C and D).

Together, these findings reveal that early NLRP3 inflammasome activation promotes macrophage-mediated control of *T. gondii* replication and suggest that GSDMD-driven pyroptosis can paradoxically limit this protective response.

### Pharmacological inhibition of GSDMD enhances host control of *T. gondii*

Given the detrimental role of GSDMD during *T. gondii* infection, we next assessed whether its pharmacological inhibition could represent a therapeutic strategy. We evaluated disulfiram (DSF), a known inhibitor of GSDMD pore oligomerization, in both in vivo and in vitro models of infection. Mice were treated intraperitoneally with vehicle or DSF 1 hour prior to infection, and splenic parasite burden was quantified four days post-infection (Figure 6A). DSF-treated animals exhibited a significant reduction in parasite burden compared to vehicle-treated controls (Figure 6B), recapitulating the resistant phenotype observed in GSDMD^−/-^ mice.

**Figure 6.**
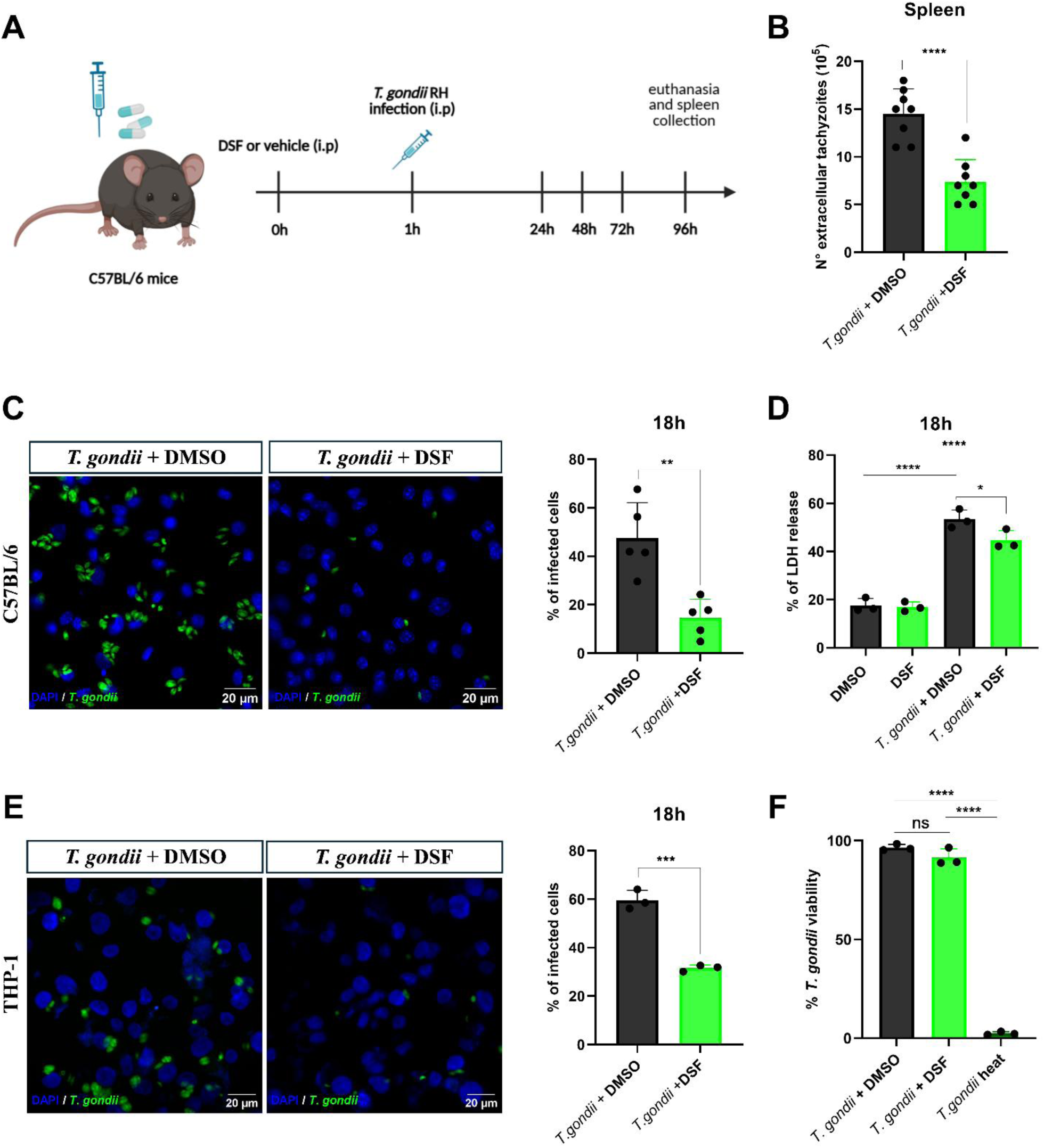
Pharmacological inhibition of GSDMD enhances host control of *T. gondii*. (A–B) C57BL/6 mice (6–10 weeks old) were pretreated with disulfiram (DSF; 50 mg/kg) or vehicle 1 h prior to intraperitoneal infection with 1 × 10⁴ *T. gondii* RH-YFP tachyzoites. (B) Four days post-infection, animals were euthanized and extracellular tachyzoites in the spleen were quantified. Data are pooled from two independent experiments (n = 8 mice per group). (C–D) Bone marrow–derived macrophages (BMDMs; 2 × 10⁵ cells/well) from C57BL/6 mice were pretreated with disulfiram (10 µM) or vehicle for 1 h and subsequently infected with *T. gondii* RH-YFP tachyzoites (MOI 2) for 18 h. After infection, culture supernatants were collected and cells were fixed with PFA and stained with DAPI. (C) Infection index was quantified by immunofluorescence microscopy. (D) Cell death was assessed by measuring LDH release in culture supernatants. (E) Human THP-1 cells were plated (1 × 10⁵ cells/well) and differentiated into macrophage-like cells with PMA (10 nM) for 24 h followed by a 24 h resting period. Cells were then pretreated with disulfiram (10 µM) for 1 h and infected with *T. gondii* RH-YFP tachyzoites (MOI 2) for 18 h. Infection index was quantified by immunofluorescence microscopy. (F) To evaluate potential direct effects of disulfiram on parasite viability, *T. gondii* tachyzoites were incubated with DMSO or DSF (10 µM) for 1 h. Heat-inactivated parasites (100 °C, 5 min) were used as a positive control. Parasite viability was assessed using the CellTiter Blue assay. Images were acquired using an IN Cell Analyzer 2200 microscope and quantified using ImageJ. Panel C represents the mean of five independent experiments performed in technical triplicates. Panel D represents the mean of three independent experiments performed in technical triplicates. Panel E shows data from one representative experiment of two independent experiments performed in technical triplicates. Panel F represents the mean of three independent experiments performed in technical sextuplicate. Statistical analysis was performed using Student’s *t*-test or one-way ANOVA. *p* < 0.05, **p** < 0.01, ***p*** < 0.001, **p** < 0.0001; ns, not significant.

Consistently, DSF treatment of BMDMs resulted in decreased intracellular parasite replication (Figure 6C) and reduced LDH release upon infection (Figure 6D), mirroring the phenotype of GSDMD-deficient macrophages. Importantly, a similar reduction in parasite replication was observed in human THP-1 derived macrophage-like cells (Figure 6E), suggesting that the protective effect of GSDMD inhibition is conserved across species.

To exclude the possibility that DSF directly compromises parasite viability, tachyzoites were treated with DSF and viability was assessed. While heat-inactivated parasites were nonviable, DSF treatment did not significantly affect tachyzoite viability compared to untreated controls (Figure 6F). Together, these findings indicate that the protective effects associated with GSDMD inhibition are unlikely to result from direct parasiticidal activity of DSF and support GSDMD as a potential therapeutic target for limiting *T. gondii* replication.

## Discussion

In this study, we demonstrate that GSDMD plays an unexpected detrimental role during infection with the highly virulent RH strain of *Toxoplasma gondii*. Both genetic deletion and pharmacological inhibition of GSDMD enhanced macrophage-mediated control of parasite replication. Mechanistically, this phenotype was associated with accelerated NLRP3 inflammasome activation and increased early IL-1β release. Together, these findings reveal that GSDMD-dependent cell death can paradoxically limit protective inflammasome responses during *T. gondii* infection.

*Toxoplasma gondii*, first described in 1908 in Tunisia and later independently identified in Brazil, is the etiological agent of toxoplasmosis (33). This globally distributed zoonotic disease is characterized by a complex epidemiology and a wide spectrum of clinical manifestations. The evolutionary success of this parasite relies on its ability to establish long-term infections in intermediate hosts while avoiding excessive host damage. Such balance imposes strong selective pressure for the development of parasite strategies that modulate host immune responses (34,35). GSDMD has been identified as a target of pathogen-mediated modulation in different infectious settings (19,36–38). Virulence factors from the RH strain of *T. gondii* have been shown to suppress GSDMD-NT activity (29), which could explain the presence of GSDMD C-terminal fragment in the supernatant of infected macrophages coupled with the absence of the N-terminal fragment, thus suggesting that regulation of this pathway may be important for host–pathogen interactions.

Inflammasome activation promotes the cleavage of GSDMD, leading to pore formation in the plasma membrane required for pyroptosis and IL-1β/IL-18 release (39). These processes contribute to the elimination of intracellular replicative niches and promote the recruitment and activation of additional immune cells (40). However, excessive or premature induction of pyroptosis can also be detrimental to the host, promoting pathological inflammation and tissue damage. Understanding how this pathway is regulated is therefore critical to balance effective pathogen control with host tissue homeostasis.

Although GSDMD has been implicated in host defense against intracellular pathogens (17,19,23,41), its role during infection with *T. gondii*, particularly with highly virulent strains such as RH, remains poorly defined. In this study, we show that while Casp1/11-deficient mice exhibit increased susceptibility to infection, GSDMD-deficient animals display the opposite phenotype, with enhanced control of parasite replication. These findings indicate that protective immune responses downstream of caspase-1 can occur independently of GSDMD. A similar phenotype has been reported in fungal infection models, in which GSDMD deficiency also resulted in improved pathogen control (21), highlighting a context-dependent role for this molecule in host responses to infection.

Consistent with the *in vivo* findings, Casp1/11^−/-^ macrophages displayed impaired control of *T. gondii* replication, whereas GSDMD-deficient macrophages exhibited enhanced resistance to infection. Importantly, no differences in infection levels were observed at early time points following parasite exposure, indicating that GSDMD deficiency does not affect parasite invasion but rather influences host–parasite interactions during later stages of infection. Enhanced parasite control in GSDMD-deficient macrophages was associated with increased IL-1β release during the early stages of infection. While Casp1/11-deficient macrophages completely lost IL-1β secretion, GSDMD^−/-^ macrophages produced significantly higher levels of this cytokine shortly after infection. The functional relevance of this early IL-1β response was further supported by the observation that exogenous IL-1β administration restored parasite control in WT macrophages to levels comparable to those observed in GSDMD-deficient cells. These observations are consistent with previous studies showing increased susceptibility to *T. gondii* infection in IL-1R-deficient mice (12), supporting a protective role for the caspase-1–IL-1β–IL-1R signaling axis in host resistance to this parasite.

Although IL-1β secretion is classically dependent on GSDMD pore formation, several alternative cytokine release mechanisms have been described. For example, infection with the type II Prugniaud strain of *T. gondii* can induce IL-1β secretion through a non-canonical pathway involving Syk–CARD9/MALT1–NF-κB signaling (42). In addition, other members of the gasdermin family have been implicated in membrane permeabilization and cytokine release. GSDMB, for instance, has been shown to mediate pore formation and pyroptosis induction (43). These alternative gasdermins may therefore contribute to membrane damage and IL-1β release during *T. gondii* infection in the absence of GSDMD. IL-1β release and cell death induction observed in GSDMD knockout cells may also be linked to NINJ1-mediated mechanisms. In the context of influenza A virus infection, IL-1β release occurs independently of GSDMD but partially depends on NINJ1 (44). In our model, NINJ1 deficiency impaired IL-1β secretion and increased parasite replication, supporting a protective role for this molecule and indicating that NINJ1-dependent membrane rupture can contribute to cytokine release independently of GSDMD activation.

Our results further indicate that enhanced resistance to *T. gondii* infection in GSDMD-deficient macrophages is associated with increased NLRP3 inflammasome activation. Immunofluorescence analysis revealed a higher number of NLRP3 puncta in GSDMD^−/-^ macrophages, suggesting increased inflammasome assembly. Increased NLRP3 activation, rather than simply prolonged cell survival, likely contributes to the elevated IL-1β response observed in these cells. Supporting this interpretation, pharmacological inhibition of NLRP3 with MCC950 partially reversed the resistant phenotype of GSDMD-deficient macrophages, indicating that early inflammasome activation is required for enhanced parasite control.

Together, these findings support a model in which GSDMD-driven early cell death limits the temporal window for inflammasome signaling and cytokine production. In the absence of GSDMD, macrophages remain viable for longer, thereby extending the temporal window for inflammasome signaling and cytokine production. Moreover, GSDMD has been shown to interact with intracellular membranes, including mitochondria and lysosomes. Cleavage of GSDMD at these sites may compromise organelle integrity and impair early microbicidal responses (45–47). For instance, GSDMD-mediated mitochondrial damage has been associated with reduced mitochondrial abundance and impaired cellular metabolism. Thus, preservation of mitochondrial function in GSDMD-deficient macrophages could contribute to improved cellular fitness and enhanced parasite control during infection. In this context, NINJ1 and GSDMD appear to exert non-redundant functions, with NINJ1 supporting cytokine release and host protection, while GSDMD-driven pyroptosis restricts the early IL-1β–dependent response.

The enhanced resistance observed in GSDMD-deficient macrophages may also involve downstream effector mechanisms of IL-1β signaling. Indoleamine 2,3-dioxygenase (IDO) and nitric oxide synthase 2 (NOS2/iNOS) are key enzymes involved in restricting *T. gondii* proliferation (48–52). However, our preliminary data indicate that GSDMD deficiency does not significantly impact IDO induction or nitric oxide production in BMDMs, suggesting that additional IL-1β–dependent effector mechanisms may contribute to enhanced parasite control.

The detrimental role of GSDMD in macrophage control of *T. gondii* infection also raises the possibility that this molecule may represent a potential therapeutic target. In agreement with this hypothesis, pharmacological inhibition of GSDMD with disulfiram improved parasite control both *in vitro* and *in vivo*, recapitulating the phenotype observed in GSDMD-deficient cells. Notably, disulfiram treatment resulted in an even greater reduction in parasite burden than genetic deletion of GSDMD. This difference may reflect the ability of disulfiram to inhibit additional gasdermin family members or interfere with metabolic pathways that are essential for parasite survival. Given that disulfiram is already FDA-approved for clinical use, our findings may accelerate its potential repurposing for the treatment of toxoplasmosis.

In summary, our findings reveal a previously unappreciated detrimental role for GSDMD during infection with virulent *T. gondii*. By promoting early inflammatory cell death, GSDMD limits protective NLRP3–IL-1β signaling and reduces macrophage-mediated control of parasite replication. These findings highlight the context-dependent functions of gasdermins during infection and suggest that modulation of GSDMD activity may represent a promising strategy to enhance host resistance to *T. gondii*.

## ACKNOWLEDGEMENTS

We thank Elisabeth de Souza for the valuable technical support.

## AUTHOR CONTRIBUTIONS

R.Q.S conceptualized the study, performed the experiments, analyzed the results, and wrote and edited the paper. A.V.C performed the experiments, analyzed the results, and reviewed the paper. L.Z.M.F.B contributed to the study’s conceptualization and reviewed the paper. M.J.D. performed THP-1 cell culture and differentiation into macrophage-like cells. K.R.B conceptualized the study, guided the data analysis, obtained funding, wrote, and reviewed the paper.

